# Efficient modular system identification provides a high-resolution assay of temporal processing and reveals the multilevel effects of attention along the human auditory pathway

**DOI:** 10.1101/2024.08.11.607503

**Authors:** Ravinderjit Singh, Hari Bharadwaj

## Abstract

Human studies of auditory temporal processing and the effects therein of aging, hearing loss, musicianship, and other auditory processing disorders have conventionally employed brainstem evoked potentials (e.g., FFRs/EFRs targeting specific modulation frequencies). Studies of temporal processing in forebrain structures are fewer and are often restricted to the 40 Hz steady-state response. One factor contributing to the limited investigation is the lack of a fast and reliable method to characterize temporal processing non-invasively in humans over a wide range of modulation frequencies. Here, we use a system-identification approach where white noise, modulated using an extended maximum-length sequence (em-seq), is employed to target stimulus energy toward a modulation-frequency range of interest and efficiently obtain a robust auditory modulation-temporal response function or ‘mod-TRF’. The mod-TRF can capture activity from sources in the early processing pathway (5-7 ms latency), middle-latency region (MLR), and late latency region (LLR). The mod-TRF is a high-resolution, modular assay of the temporal modulation transfer function (tMTF) in that the distinct neural components contributing to the tMTF can be separated on the basis of their latency, modulation frequency band, and scalp topography. This decomposition provides the insight that the seemingly random individual variation in the shape of the tMTF can be understood as arising from individual differences in the weighting and latency of similar underlying neural sources in the composite scalp response. We measured the mod-TRF under different states of attention and found a reduction in latency or enhancement of amplitude of the response from specific sources. Surprisingly, we found that attention effects can extend to the earliest parts of the processing pathway (5ms) in highly demanding tasks. Taken together, the mod-TRF is a promising tool for dissecting auditory temporal processing and obtain further insight into a variety of phenomenon such as aging, hearing loss, and neural pathology.

## 1 Introduction

Temporal processing occurs at different scales along the auditory processing pathway, with microsecond precision in the periphery to tens of milliseconds precision more centrally (JORIS et al., 2004). Currently, there is no established measure to reliably and efficiently assess temporal processing noninvasively along the entire processing pathway at the level of individual human listeners. Previous investigations in auditory science have primarily focused on characterizing early parts of the processing pathway (less than 10 ms latency) via brainstem evoked potentials (e.g., ABRs and FFRs). Investigations of more central processing areas have tended to focus on single-frequency envelope following responses (EFRs), such as the 40 Hz EFR. Single-frequency EFRs offer a limited perspective on temporal processing in individuals because their net magnitude is strongly influenced by individual anatomical variations (e.g., head sizes and tissue geometry) and the relative latencies of the multiple underlying sources that contribute to the overall response; specifically, how individual sources add constructively or destructively at the scalp electrode can obscure the true physiological variations in temporal processing (Kuwada et al., 2002; Purcell et al., 2004). Consistent with this notion, recent work by Gransier et al. (2021) shows that the temporal modulation transfer function (tMTF: a measure of phase-locked response amplitudes as a function of modulation frequencies) can be highly variable from person to person. These observations underscore the need to characterize temporal processing with a broadband approach that can separate the underlying sources contributing to the tMTF; however, measuring a full tMTF as Gransier et al. did can require up to 6 hours of recording time per participant, which renders it impractical for studying individual differences in temporal processing at a larger scale. Given that altered temporal processing is implicated in aging, hearing loss, musicianship, and neuropsychiatric disorders (Harris and Dubno, 2017; Brenner et al., 2003; Bacon and Viemeister, 1985; Gransier et al., 2020), there is a clear need to efficiently characterize temporal processing along the auditory pathway noninvasively and robustly in individuals. Here, we describe the development and validation of a system identifcation approach, yielding the modulation temporal response function (mod-TRF), to address this gap.

Measuring temporal processing in more central areas requires additional considerations compared to earlier stages of the auditory pathway (e.g., latency *<* 10 ms of processing). Not only are the responses from more central neurons sensitive to the overall state of arousal of the individual (awake vs. asleep, attending vs. passive, etc.) (Varghese et al., 2015), they also show nonlinear effects depending on the statistics of the incoming stimulus. For example, subcortical responses to speech and music converge once peripheral nonlinearities are accounted for, whereas cortical responses remain different (Shan et al., 2024). Similarly, the amplitude of the response of the cortical pyramidal-interneuron-gamma (PING) network is resonantly enhanced for stimuli with periodicities in the gamma frequency range (Fries et al., 2007). Furthermore, cortical neurons express their selectivity to preferred stimulus features more strongly when they are embedded within ongoing stimuli and are less selective at stimulus onset or for brief stimuli (Wang, 2007). Therefore, depending on the stimulus paramaters used for temporal-processing measurements, considerably different results can be obtained. Here, we sought to utilize an approach that is more akin to realistic listening conditions in that the stimulus is ongoing rather than brief, broadband rather than tonal, and aperiodic rather than regularly modulated. Another important consideration when selecting an approach is experimental efficiency. For example, recent work has shown that tMTFs between individuals can exhibit substantial variability (Gransier et al., 2021). To study temporal processing in central areas more broadly, and to elucidate the significance of this individual variability, it is crucial that measures can be obtained much more rapidly. The measures presented in this work require considerably shorter recording times compared to previous approaches of building the tMTFs from discrete modulation frequencies (Gransier et al., 2021) (of the order of 10x faster).

Broadband systems identification approaches provide an efficient method to measure temporal processing along the auditory system with an ongoing, broadband, and aperiodic stimulus. Broadband systems identification approaches using speech-like stimuli and regularized linear regression have been used to investigate cortical responses, but these approaches have poor signal-to-noise ratio (SNR) at the individual level and are studied with group averages (Crosse et al., 2016; Lalor et al., 2009; Kulasingham and Simon, 2023). An approach that uses “peaky speech” to obtain a temporal response function via deconvolution (without regularization) may potentially be useful for evaluating temporal processing; however, the approach is primarily focused on studying subcortical temporal processing currently (Polonenko and Maddox, 2021). The systems identification approach introduced here utilizes an extended maximum length sequence (em-seq) to obtain a modulation temporal response function (mod-TRF) of the auditory pathway. Crucially, the em-seq can be parametrically modified to focus the characterization energy to responses from earlier components of the auditory pathway, or to estimate the strength of responses spanning early, middle, and late latency responses simultaneously. In previous work, we described the use of em-seqs to study cortical processing of dynamic binaural cues, whereas here we focus on the more general question of amplitude modulation processing (Singh and Bharadwaj, 2023). Our results indicate that the mod-TRF, while exhibiting substantial individual differences from person to person, is highly repeatable for a given individual, enhancing the applicability of the approach to study individual physiological differences in temporal processing. By virtue of providing a broadband characterization, the mod-TRF allows for separation of the underlying sources that are emphasized in different ranges of modulation frequencies. Thus, the mod-TRF is modular, in that we can use estimates of temporal latency, modulation frequency coverage of the tMTF, and scalp topography to delineate the different sources that contribute to the overall response. Thus, the mod-TRF can be a powerful tool for assessing temporal processing in the auditory system with potential applications to hearing loss, musicianship, and neuropsychiatric diseases. Finally, because the mod-TRF includes both early and late components of neural responses, we also anticipated that they can capture differences in the arousal state of the individual. To test this, we measured the mod-TRF in both a passive state and an attentive state while performing two different tasks of varying difficulty. Neural correlates of attention are typically studied in selective-attention paradigms where participants attend to one source while ignoring another (Golumbic et al., 2013; Choi et al., 2014; O’sullivan et al., 2014; Viswanathan et al., 2019). Here, we examine the effects of attention on the mod-TRF by fixing the stimulus while manipulating the nature and level of attentional demand through different tasks. We found that there are individual differences in how attention affects the mod-TRF, as well as consistent patterns of latency and amplitude shifts in particular sources contributing to the overall mod-TRF. Surprisingly, we found that when attentional demands are high and sustained, even the short-latency (≈ 5 ms latency) component of the mod-TRF is altered. These effects were not observed when the attentional demands were weaker. In summary, the mod-TRF promises to be a tool for characterization of temporal processing with a greater level of detail and efficiency compared to conventional approaches.

## 2 Methods

### 2.1 Human Participants

Nine participants (6 female), with an average age of 26 years (ranging from 23 to 30), were recruited from the Greater Lafayette area via posted flyers and advertisements. Audiograms were measured using calibrated Sennheiser HDA 300 headphones, following a modified Hughson-Westlake procedure. All participants had hearing thresholds of 25 dB HL or better in both ears across octave frequencies from 250 Hz to 8 kHz, including 3 and 6 kHz. Informed consent was obtained from all participants, and all procedures were approved by the Institutional Review Board and the Human Research Protection Program at Purdue University.

### 2.2 EEG recording

Digital stimuli were created using custom scripts in MATLAB (The MathWorks Inc., Natick, MA), sampled at a rate of 48,828.125 Hz. These digital signals were converted to analog voltage signals using an RZ6 audio processor (Tucker-Davis Technologies, Alachua, Florida) and then delivered to the participants’ ears via ER2 insert earphones (Etymotic Research, Elk Grove Village, IL) with foam ear tips.

EEG measurements were recorded using a 32-channel Biosemi Active II system (Biosemi, Amsterdam, Netherlands). EEG data were sampled at 4096 Hz and filtered between 1-200 Hz. The data were rereferenced to electrodes placed on the earlobes. Blink artifacts were removed using the signal-space projection method (Uusitalo and Ilmoniemi, 1997). The noise-space weights were manually selected for each participant to match the topographic pattern expected for blink artifacts.Passive EEG data were collected while participants watched a muted video with subtitles of their choosing, as the stimuli were played. Active EEG data were collected by having participants respond using a button box after the stimuli were presented. Trials with peak-to-peak deflections greater than 200-300 *µ*V were excluded to minimize movement artifacts.

### 2.3 Systems Identification using extended m-seq

We utilized an extended m-sequence (em-seq) approach, previously used to study dynamic binaural processing (see Singh and Bharadwaj (2023)), and extended it here to study temporal processing, thereby obtaining a modulation temporal response function (mod-TRF). While some details are revisited here, a more complete overview of the em-seq approach can be found in Singh and Bharadwaj (2023). Fig. 1 illustrates the methodology used in this study. An em-seq modulates white noise as the stimulus. In each trial, a different realization of white noise is paired with the same em-seq. There are two parameters that need to be specified when designing an em-seq: the number of bits, *n*, and f_4*dB*_, the frequency at which the em-seq loses approximately 4 dB of energy. The number of bits controls the length of the em-seq. While employing longer sequences (higher *n*) offers an SNR benefit, the minimum technical requirement is that the signal length must exceed the duration of the system function being measured. The choice of f_4*dB*_ also determines the cut-off frequency (COF = 2 * f_4*dB*_) and the duration of each point in the m-seq (T = 1/COF) to obtain the em-seq. F_4*dB*_ serves as a boundary for characterizing the system’s response. The em-seq loses 4 dB in power at f_4*dB*_ and then rapidly loses the rest of its energy between f_4*dB*_ and the COF. For example, if f_4*dB*_ is 100 Hz, the COF will be 200 Hz, and T will be 5 ms. Using 9 bits, the underlying m-seq has a length of 2^9^ − 1, and with each point being T = 5 ms, the total duration is 2.56 seconds.

**Figure 1.**
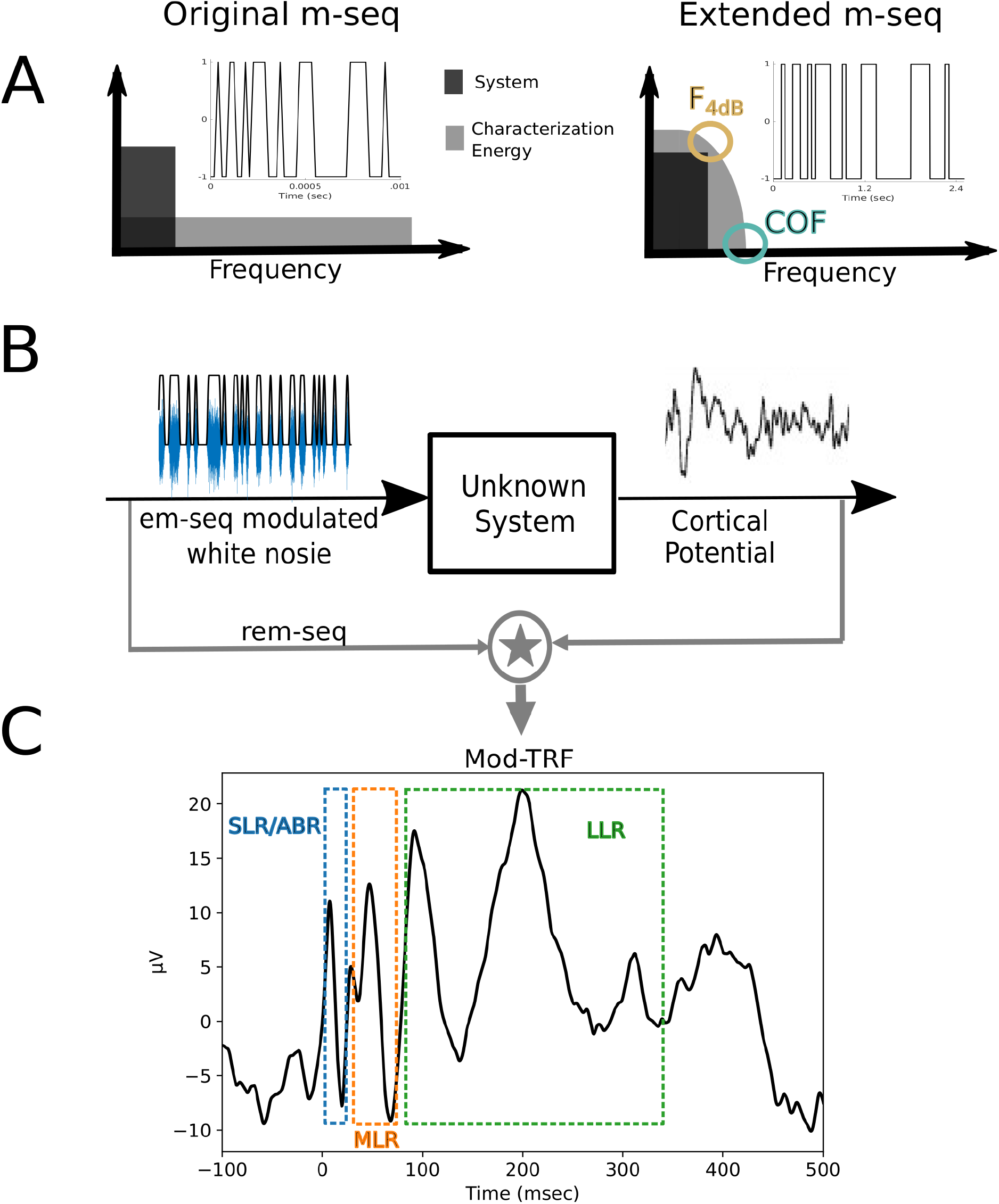
**A** illustrates the transformation of the m-seq into the extended m-seq by increasing the duration of each point in the m-seq. This transformation modifies the frequency response of the m-seq to a sinc shape, rather than white, which helps focus the characterization energy within the range where the system of interest is active. f_4*dB*_ denotes the frequency at which 4 dB of characterization energy is lost, and the cut-off frequency (COF) at 2 * f_4*dB*_ is where characterization energy reaches zero. **B** depicts the paradigm used to obtain system responses. The extended m-seq modulated white noise is played while voltage potentials (EEG) are recorded. The EEG signals from each channel are cross-correlated with the recovery em-seq to estimate the system response. **C** shows an example system response (mod-TRF) obtained in this study, highlighting the SLR, MLR, and LLR regions, which are captured with f_4*dB*_ = 75 Hz.

To prevent energy from spreading above our COF, the rise and fall of the em-seq were ramped. The ramp was 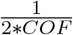 in duration, achieved by convolving the em-seq with a 1st order discrete prolate spheroidal sequence (DPSS) window of 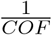 in duration, and then ensuring the em-seq’s magnitude remained between 0 and 1. To obtain the system response (mod-TRF), each EEG channel was cross-correlated with the recovery em-seq, which is the same as the em-seq but oscillates between -1 and 1. This adjustment eliminates the introduction of a DC component to the system function.

In this study, f_4*dB*_ was set at 75 Hz to obtain a good representation of late processing areas that primarily encode low-frequency modulations (JORIS et al., 2004). For better representation of earlier processing areas, a higher f_4*dB*_, between 250-500 Hz, might be more appropriate, though this comes at the cost of less energy for later processing areas, making peaks in these regions less evident. The temporal resolution of the system response is the duration of each point in the em-seq, approximately 7 ms in this case. The em-seq had 10 bits, resulting in a stimulus duration of 6.82 seconds. We collected data at 7, 8, 9, and 10 bits in 5 participants to demonstrate the change in SNR with increasing bit length. The durations for these bit lengths were 0.85, 1.70, 3.40, and 6.81 seconds, respectively. Each bit value was tested in 300 trials, resulting in total recording times of approximately 7, 11.3, 19.8, and 36.8 minutes, respectively, accounting for a 0.5-second interstimulus interval and 0.1-second random jitter between trials. These recording times are significantly shorter than the 6 hours required to measure a comprehensive tMTF (Gransier et al., 2021). Noise floors were computed by randomly inverting the polarity of half the trials and processing these noise responses through the same analysis pipeline as the actual data. Fifty noise floors were computed per participant. Fig. 2 shows the mod-TRF measured with different bit values, illustrating the improvement in SNR with increasing bit numbers. The 10-bit em-seq was measured in all 9 participants and repeated in 6 participants months apart from the initial measurement.

**Figure 2.**
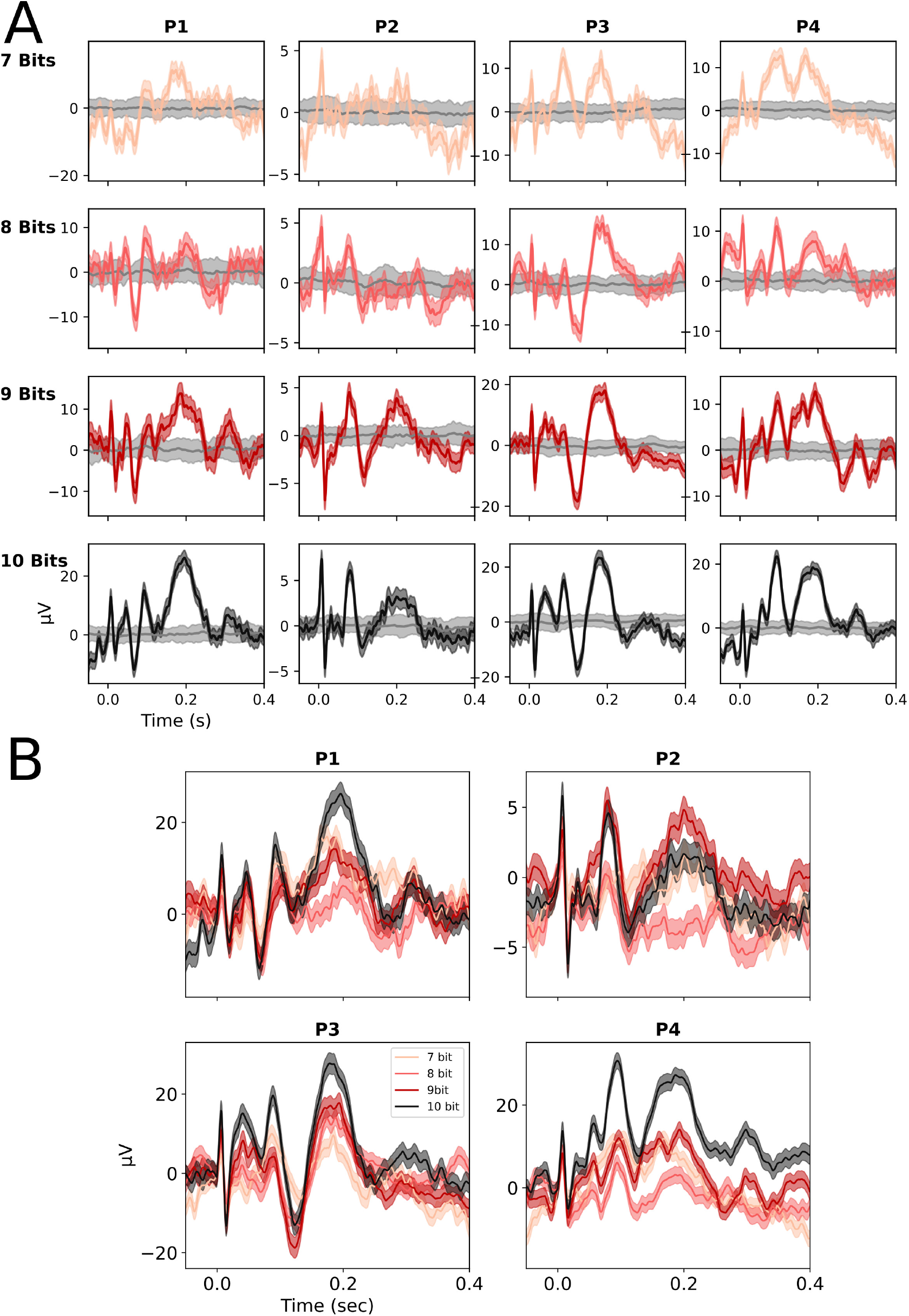
**A** The mod-TRF measured with different bit values across 4 participants. The gray line represents the mean of the noise floors, with gray shading indicating **+/- 1 standard deviation**. The colored shading represents the standard error across trials. **B** The mod-TRF measured with different bit values, with time zero set to start from 0 *µV*. Note that while the mod-TRFs are quite similar, the 9- and 10-bit measurements appear smoother and better capture the later peaks. This improvement is due to the higher SNR of larger bit value mod-TRFs and the longer duration of the mod-TRF stimuli, which contain more low-frequency energy, enhancing the characterization of the later peaks.

### 2.4 Identifying the underlying sources

A source here refers to a group of neurons firing with similar latencies and synchronously enough to form a distinguishable peak in the mod-TRF. The resolvability of distinct sources depends on the parameters of the em-seq used to obtain the mod-TRF. For instance, with f_4*dB*_ = 75 Hz, several peaks typically observed in the short latency region (i.e., ABR) with clicks are represented by a single peak, as illustrated in Fig. 4. A higher f_4*dB*_ may allow for better resolution of multiple peaks in the short latency region; however, this could also lead to energy available for clearly identifying sources contributing to the late latency region. Due to the limitations of recording EEG with only 32 electrodes, we did not conduct a detailed source analysis by solving the inverse problem to pinpoint the exact brain regions responsible for the activity. Instead, we aimed to categorize sources as subcortical, cortical, or mixed/indeterminate, based on peak latency, peak width (or coverage of the tMTF), and the distribution of activity across the sensors. It is well documented that the latency of each subsequent source in the auditory system increases along the ascending processing pathway. Additionally, later processing areas exhibit a progressive conversion to a rate code, or longer integration times, resulting in broader peaks and a lower pass coverage of the tMTF. To identify the different sources, we first manually marked the troughs in the mod-TRF averaged across all 9 participants and analyzed the peak latency and width of the identified peaks. We then utilized a PCA implementation from scikit-learn in Python to examine the topographic distribution of the signal between troughs by plotting the PCA weights for each component across the 32 scalp sensors. The polarity of each component was forced to be positive at channel Cz, as the sign of a PCA component is arbitrary. We plotted the PCA weights on a scalp topographic map and report the explained variance for the principal component representing each source. The explained variance serves as a proxy for how accurately the depicted topomap reflects the distribution of activity across electrodes corresponding to the underlying source. A lower explained variance indicates that the topomap is a less reliable representation of the dominant source. A source with a very broad distribution across all sensors suggests a subcortical origin, while a source with focused activity around channel Cz is more likely to originate from the bilateral auditory cortices (Scherg and Von Cramon, 1986; Cohen and Cuffin, 1983). From this analysis, we identified 5 distinct sources in the mod-TRF, as shown in Fig. 4, and subsequently manually identified these 5 sources in individual participants. In Fig. 4, the topomap for source I is consistent with a subcortical origin, while the topomaps for sources IV and V align with expected activity from the bilateral auditory cortices.

### 2.5 Active Paradigms

The passive paradigm involved participants watching a muted video with subtitles and ignoring the presented auditory stimuli, alonging with EEG recording. Two active paradigms were employed to assess how attention would alter the mod-TRF, with EEG recordings obtained during both tasks. The paradigms were labeled as ‘easy’ and ‘hard.’

In the easy paradigm, participants were instructed to count the number of times the stimulus (a 10-bit em-seq modulated white noise with f_4*dB*_ = 75 Hz) repeated. The stimulus could repeat 1-3 times, and participants responded by indicating the number of repetitions on a button box.

The hard paradigm involved a 3-alternative-forced-choice (3AFC) task, where the target interval had the em-seq modulation circularly time-shifted by 50%. While the spectral properties of the em-seq remain unchanged by circular shifting, the temporal modulation pattern becomes differentt. Participants had to detect this shifted em-seq, requiring them to pay close attention to the stimulus and listen for differences in the modulation pattern. The behavioral accuracy rates were 99% and 61% (chance = 33%), respectively, consistent with the labeling of ‘easy’ and ‘hard’ tasks. All 9 participants completed the easy task, while 7 participants undertook the hard task.

### 2.6 Spectral characteristics

We computed the fast Fourier transform (FFT) of the mod-TRF using the implementation available in the NumPy package in Python. The amplitude was normalized for the length of the transform to plot the results in *µV*. We calculated the phase response for each individual, and the average phase response is depicted in Fig. **??**. To compute the group delay, we fitted a line between 10-20 Hz and 30-70 Hz, taking the negative of the fitted slope and dividing it by 2*π*. These frequency ranges were chosen to examine the group delay before and after it undergoes significant changes between 20-30 Hz.

## 3 Results

Fig. 1 illustrates the procedure for obtaining a mod-TRF, along with an example mod-TRF. The parameters for the em-seq used in this study (f_4*dB*_ = 75 Hz) allowed us to capture the SLR/ABR, MLR, and LLR regions of auditory processing. A notable aspect of this approach is the high SNR obtained in individual participants. Fig. 3 shows two mod-TRFs from 6 participants, measured months apart. The consistency of these measurements demonstrates that the mod-TRF can be reliably measured with similar results even when taken months apart, although there is substantial inter-participant variability. Fig. 4 presents the mod-TRF averaged across 9 participants, along with PCA weight topomaps indicating the distribution of activity across the 32 sensors on the scalp for the 5 identified sources. Based on the distribution of activity, peak latency, and peak width, we determined there were 5 separable sources in the mod-TRF with f_4*dB*_ = 75 Hz. Source I appears to be subcortical, while sources IV and V appear to be cortical, as indicated by their peak latency, peak width, and the high explained variance of the PCA component with topomaps consistent with subcortical and cortical sources, respectively. Source II’s topomap resembles that of a cortical source but has a lower explained variance. Source III’s topomap does not clearly indicate whether it is subcortical or cortical. Therefore, we believe sources II and III may represent a mix of subcortical and cortical activity or are indeterminate from our analysis. A more detailed source analysis with a larger number of electrodes or a higher f_4*dB*_ may be necessary to elucidate their precise locations.

**Figure 3.**
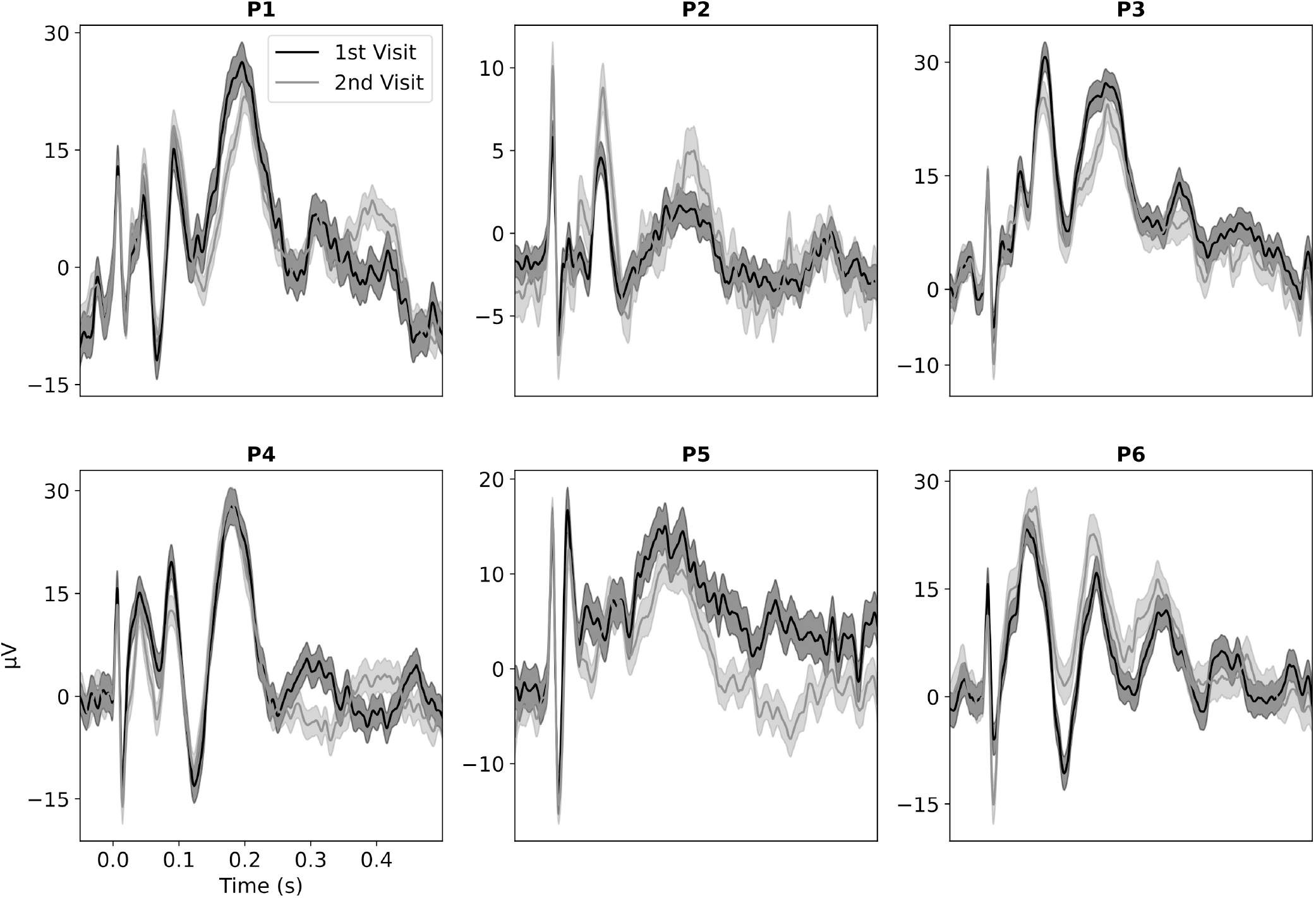
The mod-TRF measured in 6 participants across 2 visits, spaced months apart. The data suggest that the mod-TRF is repeatable with high consistency within individual participants, while also exhibiting significant variability across different participants.

**Figure 4.**
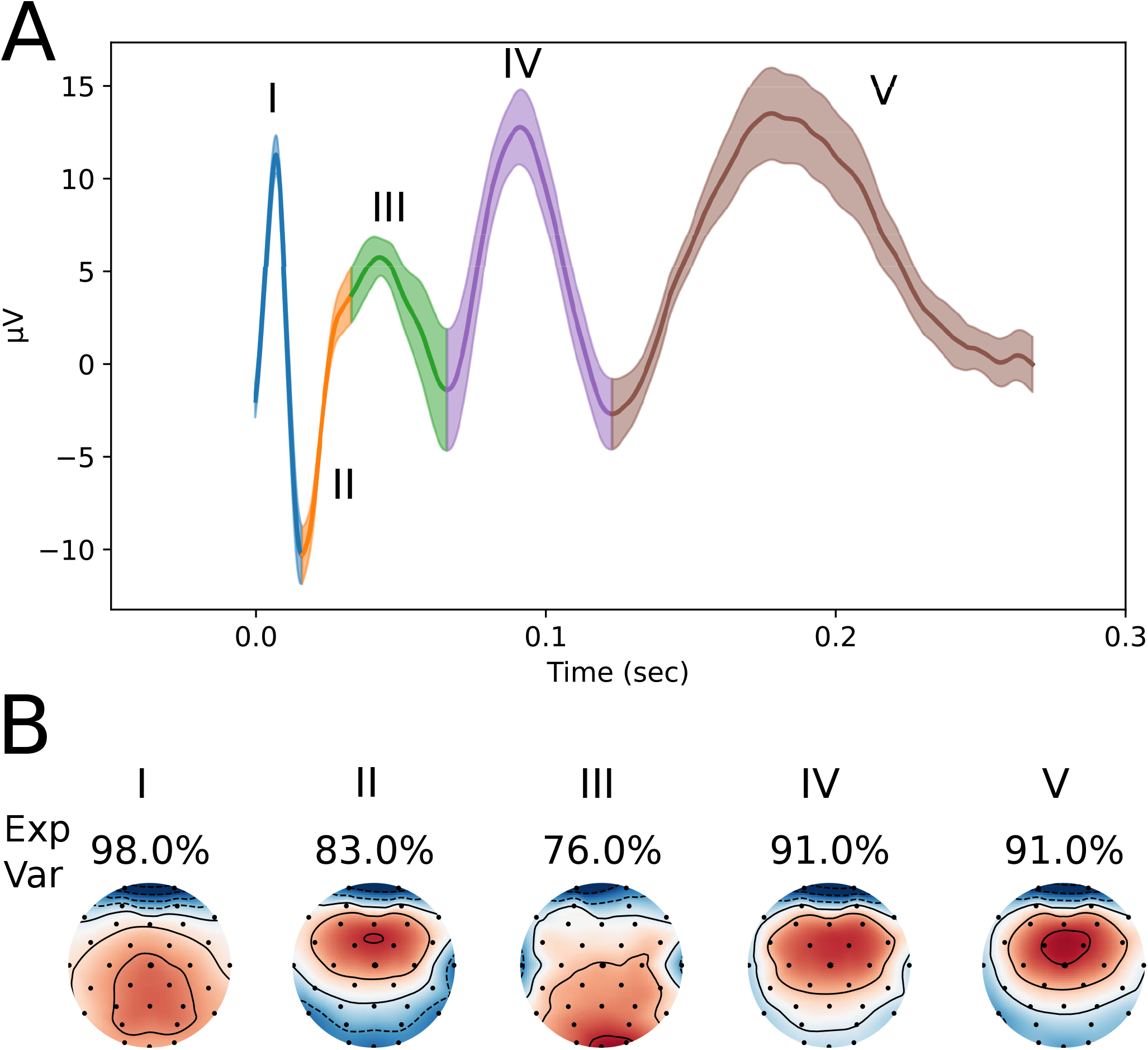
**A** The mod-TRF averaged across 9 participants in a passive state. The shading indicates the standard error, with different colors representing different sources, which are labeled with Roman numerals. **B** The PCA weights for each source plotted over topographic maps to illustrate the distribution of activity across the sensors. The amount of variance explained by each PCA component is also shown.

We separately evaluated the modulation encoding properties of each of the 5 sources, as shown in Fig. 5. The frequency response of the composite (i.e., overall) mod-TRF displays several peaks and nulls, similar to the findings of Gransier et al. when evaluating tMTFs across individuals using discrete modulation-frequency stimuli (Gransier et al., 2021). However, examining the frequency coding of individual sources reveals more low-pass characteristics, as might be expected from single neuron recordings (JORIS et al., 2004). Thus, focusing on the modulation coding of individual sources provides a more readily interpretable representation of the underlying physiological temporal coding characteristics than the frequency response of the entire mod-TRF, which is influenced by the superposition of multiple source responses at the scalp. Fig. **??** shows the average frequency response across 9 participants. After averaging across participants, many individual peaks and nulls are averaged out, resulting in a mostly low-pass response, as anticipated. The average phase response is similar to that measured by Gransier et al., showing an abrupt change in group delay (slope of the phase response) for modulation frequencies 20 Hz (Gransier et al., 2021). The average frequency response diverges from responses measured by single-frequency EFRs in the gamma frequency range (30-60 Hz). Typically, there is a strong response to periodic modulations in the gamma frequency range with the response peaking around the classic ASSR modulation frequency of 40 Hz; however, we measured the tMTF with an aperiodic stimulus, which likely does not activate circuits in the brain with resonances in the gamma frequency range, such as the PING network (Fries et al., 2007). Consistent with this notion, we do not observe strong activity in the gamma frequency range when measuring the tMTF with em-seqs.

**Figure 5.**
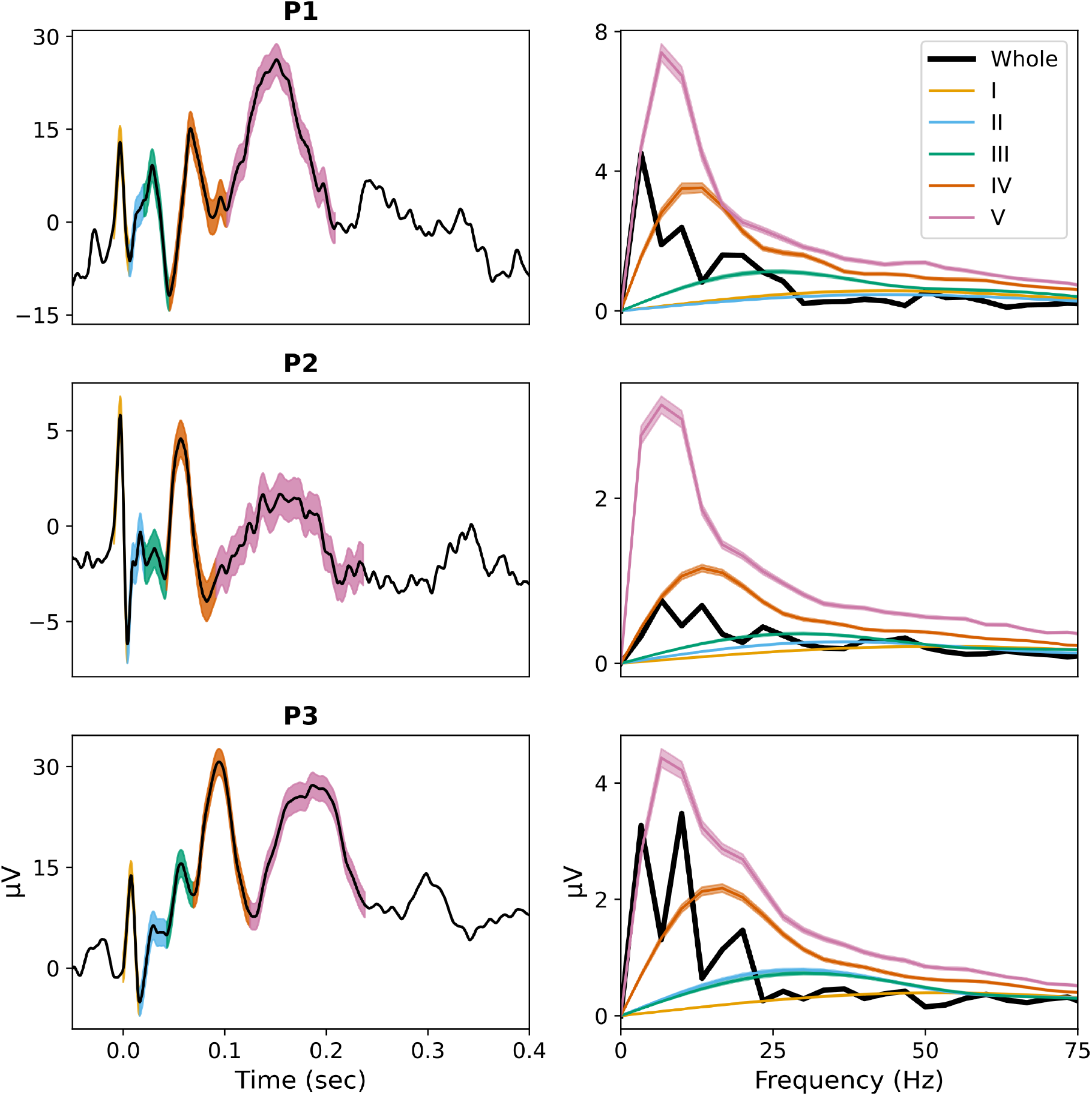
On the left is the time-domain mod-TRF, and on the right is the fast Fourier transform. The shading represents the standard error, with the color indicating a specific source. Note that the frequency domain representation of the entire mod-TRF (in black) displays several peaks and troughs, whereas the modulation coding properties of each source exhibit a low-pass/band-pass shape.

**Figure 6.**
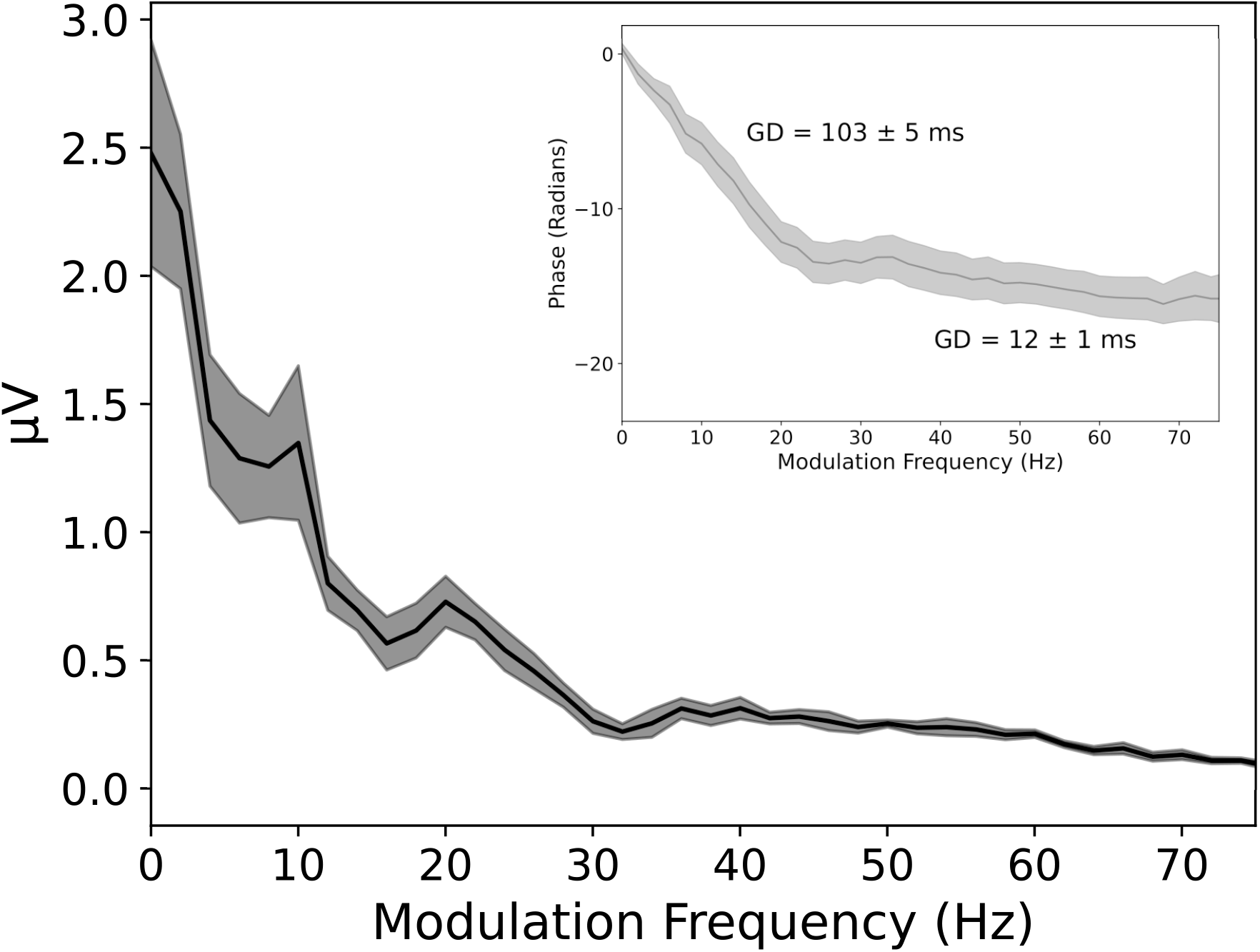
The average frequency response of the entire mod-TRF across 9 participants in the passive state. The gray line represents the phase response. The group delay, computed between 10-20 Hz and 40-60 Hz, is also shown along with the standard error.

By virtue of disentangling the multiple sources contributing to modulation coding, the mod-TRF is also a valuable tool for evaluating how attention affects each individual source. To explore this, we measured the mod-TRF under two different active tasks. The ‘easy’ task involved counting the number of times the stimulus repeated (average accuracy = 99%), while the ‘hard’ task required participants to detect a shift in the em-seq modulation (average accuracy = 61%, with chance = 33%). Changes in individual mod-TRFs for four participants under these two active conditions are shown in Fig. 7. There are individual differences in how attention affects source amplitude and latency, and it appears the two different active tasks impact sources differently. Fig. 8 presents the averaged response to the passive and active conditions, along with topomaps derived from our PCA analysis. In both active tasks, there appears to be a reduction in latency in source V and a reduction in latency and increase in amplitude of source II. Additionally, the increase in activity from source II appears subcortical in nature, according to the associated topomap distribution. However, there seems to be enhanced activity in posterior electrodes not typically seen in a standard subcortical topomap (source I has a typical subcortical topomap).

**Figure 7.**
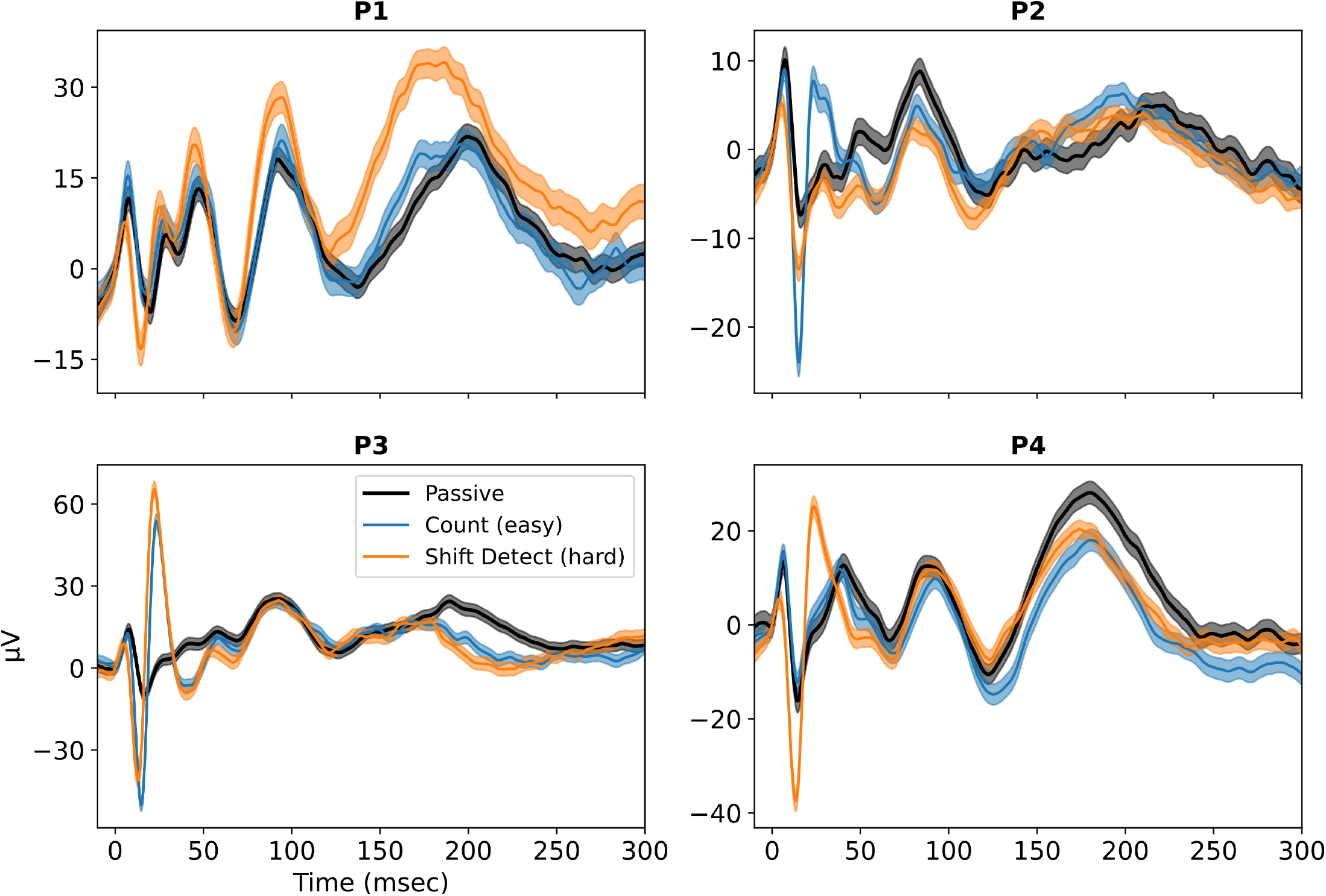
The mod-TRF measured in the passive and 2 active conditions in 4 participants from channel Cz. Note the individual differences in changes to the mod-TRF with attention.

**Figure 8.**
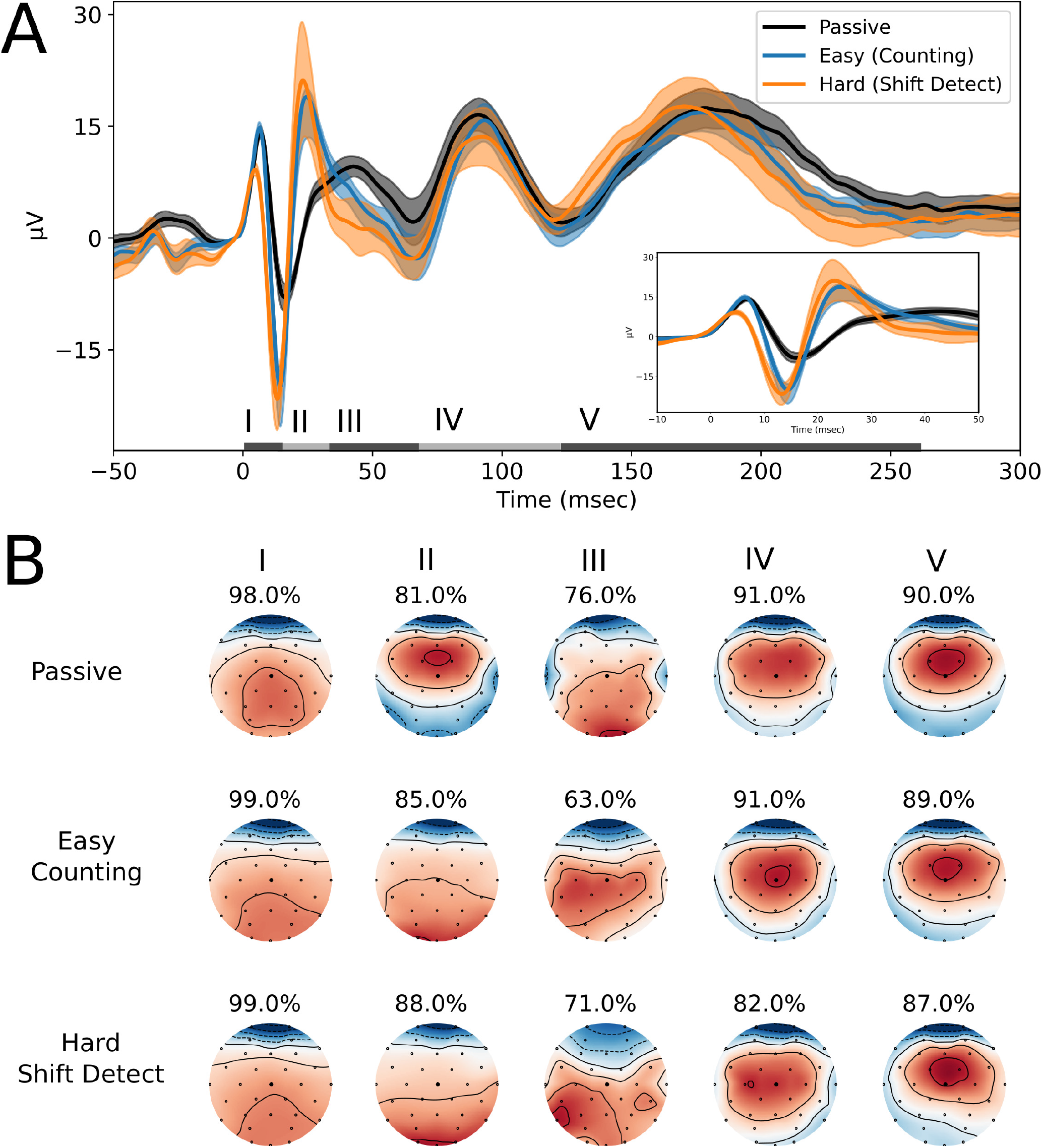
**A** Averaged responses for the mod-TRF in the passive condition (n=9), easy active task (n=9), and hard active task (n=7) from channel Cz. The shading represents the standard error, and sources are annotated at the bottom. **B** Topographic maps obtained with PCA for each source in the passive, easy counting task, and hard shift detection task conditions. The percentages above the topomaps represent the explained variance of the source.

The peak in source I, located in the SLR/ABR region, is only altered in the hard task, not the easy task, as seen in Fig. 8. It is surprising that the effects of attention extend to such early (low-latency) portions of the auditory processing pathway. However, there is some evidence that top-down modulation of the the ABR is possible in that electrical cortical stimulation modulted the ABR (Liu et al., 2019). Because the em-seq we employed only contained energy up to about 75 Hz, it would be useful our replicate this experiment with f_4*dB*_ = 250-500 Hz or higher to obtain a better resolve SLR and MLR activity, and further understand the nature of the strong effects of attention that are apparent selectively in the hard task.

## 4 Discussion

Our goal was to develop a method to rapidly assess temporal processing, and do so in a modular fashion, where the characterization energy could be parametrically focused on different ranges of modulation frequencies. To achieve this, we utilized a white noise stimulus modulated by an em-seq to measure a mod-TRF. This approach offers flexibility, allowing researchers to focus on SLR, MLR, or LLR regions, or to obtain a representation of all three, as was done in this study. The high SNR and repeatability of the observed in individuals makes the mod-TRF a promising tool for studying individual differences across various populations, such as those with hearing loss, neuropsychiatric disorders, aging, and more. The broadband nature of the input modulation energy allows for the separation of different sources contributing to the response, providing a more readily interpretable representation of the underlying physiological temporal coding processes compared to narrowband approaches (e.g., single-frequency EFR). The ability to separate sources also facilitates interesting studies on auditory physiology, such as evaluating how attention affects different parts of the auditory system. We believe the mod-TRF has potential for a wide-range of applications in studying auditory temporal processing and activity at the level of individual listeners. Additionally, this approach could be easily adapted to study temporal processing in other sensory modalities.

### 4.1 Mod-TRF and individual differences

Systems identification approaches have a long history of use in neuroscience (Eggermont, 1993; Ringach and Shapley, 2004; Recio-Spinoso et al., 2005; Lalor et al., 2006, 2009; Crosse et al., 2016; Maddox and Lee, 2018; Polonenko and Maddox, 2021). Current approaches using scalp EEG, while robust at the group level, often suffer from poor SNR at the individual level. However, studying group-averaged responses has still provided valuable insights (Ding and Simon, 2013; O’Sullivan et al., 2015). Recent work using a “peaky” speech stimulus has shown potential for obtaining robust individual responses; however, this work has focused on characterizing very early responses (less than 10 ms) (Polonenko and Maddox, 2021). We are not aware of any existing approaches the allow for the comparison of individual differences in both early and late areas of the auditory pathway.

The mod-TRF, with its high SNR in individuals, offers the opportunity to investigate individual differences. It has been shown to be repeatable in individuals and exhibits substantial inter-participant variability, as seen in Fig. 3. The significance of this inter-participant variability in explaining behavior and alterations due to pathology presents an interesting avenue for future exploration. However, a critical question to consider is the extent to which this variability is due to physiological differences versus anatomical/geometric differences. It is crucial to separate out anatomical/geometric effects to compare physiology across individuals accurately. Why are some sources clearer in one person but not another? Why do some individuals exhibit more peaks or peaks at substantially different latencies than others? To uncover the underlying physiology, we need to compare the same underlying neural source across individuals.

Scalp EEG responses are contributed to by the entire brain. We can differentiate responses from different groups of neurons based on our understanding of latencies, temporal processing abilities, and the anatomical locations of the responding neurons. While it is likely safe to assume that most individuals have very similar anatomy (e.g., everyone has an auditory nerve, cochlear nucleus, superior olivary complex, etc., organized in a similar structure), it is less clear if we can assume other aspects of processing, such as synchrony between groups of neurons in different sources, are similar. Slight differences in geometry (such as head size) could result in interactions on the scalp being destructive between two sources in one person while being constructive in another. While the mod-TRF takes substantial steps toward parsing out some of these interactions by allowing for source separation to some extent, the limits and utility of the mod-TRF in studying individual differences remain to be fully determined. Understanding how much of the variability in the mod-TRF is due to geometry/anatomy versus physiology will be crucial for determining the next steps in uncovering and comparing the true underlying physiology across individuals.

Another key consideration in source separability using the mod-TRF is the choice of f_4*dB*_. The f_4*dB*_ parameter determines the frequency up to which the majority of the input energy lies and the temporal resolution of the mod-TRF. With a lower f_4*dB*_, more sources with similar latencies may be grouped into one peak. For example, with f_4*dB*_ set to 75 Hz in this study, the five peaks typically seen in a traditional ABR appear to be represented by just one peak. Depending on the specific research question and region of interest, the f_4*dB*_ parameter will need to be adjusted accordingly. However, f_4*dB*_ set to 75 Hz, as done here, seems to provide a representation of early, middle, and late regions, making it a suitable choice for initial investigations. A lower f_4*dB*_ would make lower latency peaks less distinct and higher latency peaks more pronounced, while a higher f_4*dB*_ would have the opposite effect.

### 4.2 The mod-TRF and attention

The mod-TRF can capture activity from early, middle, and late latency components of the response and the corresponding sources along the auditory pathway in a separable fashion. This capability allows for investigating various exciting avenues, such as evaluating how attention affects different levels of of the auditory processing hierarchy. We chose to evaluate attention both to further validate the mod-TRF and to gain insights into how attention influences different parts of the auditory system. We hypothesized that if we could measure separable sources, then attention could affect these sources independently, which in in line with our observations. Perhaps more intriguing are the individual differences in how attention impacts different sources. Due to the small number of participants in this initial investigation, we did not explore how these differences in attention effects relate to behavior, but this is an interesting avenue for future research.

Our approach to studying attention using the mod-TRF stimulus differs significantly from previous attention studies. Typically, attention studies have participants ignore one stimulus while selectively attending to another and then evaluate how the focus of attention alters the response to the attended stimulus (Viswanathan et al., 2019; Choi et al., 2013). In the present study, we varied the level of attention to a single stimulus by employing different tasks. The target stimulus allowed us to derive a mod-TRF, enabling us to observe how different sources were altered by attention. The average changes indicated task-dependent effects of attention, suggesting several avenues for further exploration of how attention impacts the auditory system. Studying attention with the mod-TRF allows for evaluation of attention effects at multiple points in the auditory system. A very exciting avenue of future work will be exploring weather individual differences in both qualitative (e.g., which sources are altered) and quantitative (e.g., the strength of the attentional modulation of a given source) aspects of the attention effects can explain individual differences in behavior.

In conclusion, we have developed and validated an efficient modular systems identification approach to measure temporal processing with high resolution at the individual level. This approach is efficient, requiring only about 30 minutes of recording time to obtain a representation of temporal processing across a wide range of frequencies. The method allows for the separation of neural sources contributing to the response and is reliable, as measurements are repeatable even months apart. Interestingly, there is notable inter-participant variability in responses. We demonstrated the application of the mod-TRF in studying attention and found that attention effects can vary based on the task at hand. The mod-TRF shows promise as a ubiquitously applicable assay for probing auditory temporal processing for a range of purposes.

## Code and Data Availability Statement

The code to reproduce the figures in this work is publicly available on GitHub (https://github.com/Ravinderjit-S/ModulationTemporalResponseFunction) and archived using Zenodo (Singh and Bharadwaj, 2024a). The EEG data is openly accessible and archived using Zenodo (Singh and Bharadwaj, 2024b).

## References

Bacon, S. P. and Viemeister, N. F. (1985). Temporal Modulation Transfer Functions in Normal-Hearing and Hearing-Impaired Listeners. Audiology, 24(2):117–134.

Brenner, C. A., Sporns, O., Lysaker, P. H., and O’Donnell, B. F. (2003). EEG Synchronization to Modulated Auditory Tones in Schizophrenia, Schizoaffective Disorder, and Schizotypal Personality Disorder. American Journal of Psychiatry, 160(12):2238–2240.

Choi, I., Rajaram, S., Varghese, L., and Shinn-Cunningham, B. (2013). Quantifying attentional modulation of auditory-evoked cortical responses from single-trial electroencephalography.

Choi, I., Wang, L., Bharadwaj, H., and Shinn-Cunningham, B. (2014). Individual differences in attentional modulation of cortical responses correlate with selective attention performance. Hearing Research, 314:10–19.

Cohen, D. and Cuffin, B. (1983). Demonstration of useful differences between magnetoencephalogram and electroencephalogram. Electroencephalography and Clinical Neurophysiology, 56(1):38–51.

Crosse, M. J., Di Liberto, G. M., Bednar, A., and Lalor, E. C. (2016). The Multivariate Temporal Response Function (mTRF) Toolbox: A MATLAB Toolbox for Relating Neural Signals to Continuous Stimuli.

Ding, N. and Simon, J. Z. (2013). Adaptive temporal encoding leads to a background-insensitive cortical representation of speech. Journal of Neuroscience, 33(13):5728–5735.

Eggermont, J. J. (1993). Wiener and Volterra analyses applied to the auditory system. Hearing Research, 66(2):177–201.

Fries, P., Nikolić, D., and Singer, W. (2007). The gamma cycle. Trends in Neurosciences, 30(7):309–316.

Golumbic, E. M. Z., Ding, N., Bickel, S., Lakatos, P., Schevon, C. A., McKhann, G. M., Goodman, R. R., Emerson, R., Mehta, A. D., and Simon, J. Z. (2013). Mechanisms underlying selective neuronal tracking of attended speech at a “cocktail party”. Neuron, 77(5):980–991.

Gransier, R., Carlyon, R. P., and Wouters, J. (2020). Electrophysiological assessment of temporal envelope processing in cochlear implant users. Scientific Reports, 10(1):15406.

Gransier, R., Hofmann, M., van Wieringen, A., and Wouters, J. (2021). Stimulus-evoked phase-locked activity along the human auditory pathway strongly varies across individuals. Scientific Reports, 11(1):143.

Harris, K. C. and Dubno, J. R. (2017). Age-related deficits in auditory temporal processing: unique contributions of neural dyssynchrony and slowed neuronal processing. Neurobiology of Aging, 53:150– 158.

Joris, P. X., Schreiner, C. E., and Rees, A. (2004). Neural Processing of Amplitude-Modulated Sounds. Physiological Reviews, 84(2):541–577.

Kulasingham, J. P. and Simon, J. Z. (2023). Algorithms for Estimating Time-Locked Neural Response Components in Cortical Processing of Continuous Speech. IEEE Transactions on Biomedical Engineering, 70(1):88–96.

Kuwada, S., Anderson, J. S., Batra, R., Fitzpatrick, D. C., Teissier, N., and D’Angelo, W. R. (2002). Sources of the scalp-recorded amplitude-modulation following response. Journal of the American Academy of Audiology, 13(4):188–204.

Lalor, E. C., Pearlmutter, B. A., Reilly, R. B., McDarby, G., and Foxe, J. J. (2006). The VESPA: a method for the rapid estimation of a visual evoked potential. Neuroimage, 32(4):1549–1561.

Lalor, E. C., Power, A. J., Reilly, R. B., and Foxe, J. J. (2009). Resolving Precise Temporal Processing Properties of the Auditory System Using Continuous Stimuli. Journal of Neurophysiology, 102(1):349– 359.

Liu, X., Zhang, O., Chen, A., Hu, K., Ehret, G., and Yan, J. (2019). Corticofugal Augmentation of the Auditory Brainstem Response With Respect to Cortical Preference. Frontiers in systems neuroscience, 13:39.

Maddox, R. K. and Lee, A. K. C. (2018). Auditory Brainstem Responses to Continuous Natural Speech in Human Listeners. eNeuro, 5(1):ENEURO.0441–17.2018.

O’sullivan, J. A., Power, A. J., Mesgarani, N., Rajaram, S., Foxe, J. J., Shinn-Cunningham, B. G., Slaney, M., Shamma, S. A., and Lalor, E. C. (2014). Attentional selection in a cocktail party environment can be decoded from single-trial EEG. Cerebral Cortex, 25(7):1697–1706.

O’Sullivan, J. A., Shamma, S. A., and Lalor, E. C. (2015). Evidence for Neural Computations of Temporal Coherence in an Auditory Scene and Their Enhancement during Active Listening. Journal of Neuroscience, 35(18):7256–63.

Polonenko, M. J. and Maddox, R. K. (2021). Exposing distinct subcortical components of the auditory brainstem response evoked by continuous naturalistic speech. eLife, 10:e62329.

Purcell, D. W., John, S. M., Schneider, B. A., and Picton, T. W. (2004). Human temporal auditory acuity as assessed by envelope following responses. The Journal of the Acoustical Society of America, 116(6):3581–3593.

Recio-Spinoso, A., Temchin, A. N., van Dijk, P., Fan, Y.-H., and Ruggero, M. A. (2005). Wienerkernel analysis of responses to noise of chinchilla auditory-nerve fibers. Journal of neurophysiology, 93(6):3615–3634.

Ringach, D. and Shapley, R. (2004). Reverse correlation in neurophysiology. Cognitive Science, 28(2):147– 166.

Scherg, M. and Von Cramon, D. (1986). Evoked dipole source potentials of the human auditory cortex. Electroencephalography and Clinical Neurophysiology/Evoked Potentials Section, 65(5):344–360.

Shan, T., Cappelloni, M. S., and Maddox, R. K. (2024). Subcortical responses to music and speech are alike while cortical responses diverge. Scientific Reports, 14(1):789.

Singh, R. and Bharadwaj, H. (2024a). Modulation Temporal Response Function (mod-TRF) code. 10.5281/zenodo.11201331.

Singh, R. and Bharadwaj, H. (2024b). Modulation Temporal Response Function (mod-TRF) data. 10.5281/zenodo.11201229.

Singh, R. and Bharadwaj, H. M. (2023). Cortical temporal integration can account for limits of temporal perception: investigations in the binaural system. Communications Biology, 6(1):981.

Uusitalo, M. A. and Ilmoniemi, R. J. (1997). Communication Signal-space projection method for separating MEG or EEG into components. Technical report.

Varghese, L., Bharadwaj, H. M., and Shinn-Cunningham, B. G. (2015). Evidence against attentional state modulating scalp-recorded auditory brainstem steady-state responses. Brain Research, 1626:146–164.

Viswanathan, V., Bharadwaj, H. M., and Shinn-Cunningham, B. G. (2019). Electroencephalographic signatures of the neural representation of speech during selective attention. eNeuro.

Wang, X. (2007). Neural coding strategies in auditory cortex. Hearing Research, 229(1):81–93.

